# Stimulus-evoked and resting-state alpha oscillations show a linked dependence on patterned visual experience for development

**DOI:** 10.1101/2023.03.08.531549

**Authors:** Rashi Pant, José Ossandón, Liesa Stange, Idris Shareef, Ramesh Kekunnaya, Brigitte Röder

## Abstract

Persistent visual impairments after congenital blindness due to dense bilateral cataracts have been attributed to altered visual cortex development within a sensitive period. Occipital alpha (8-14 Hz) oscillations were found to be reduced after congenital cataract reversal during visual motion tasks. However, it has been unclear whether reduced alpha oscillations were task-specific, or linked to impaired visual behavior in cataract-reversed individuals. Here, we compared resting-state and stimulus-evoked alpha activity between individuals who had been treated for dense bilateral congenital cataracts (CC, n = 13, mean duration of blindness = 11.0 years) and age-matched, normally sighted individuals (SC, n = 13). We employed the visual impulse response function, adapted from VanRullen and MacDonald (2012), to test for the characteristic alpha response to visual white noise. Participants observed white noise stimuli changing in luminance with equal power at frequencies between 0-30 Hz. Compared to SC individuals, CC individuals demonstrated a reduced likelihood of exhibiting an evoked alpha response. Moreover, stimulus-evoked alpha power was reduced and correlated with a corresponding reduction of resting-state alpha power in the same CC individuals. Finally, CC individuals with an above-threshold evoked alpha peak had better visual acuity than CC individual without an evoked alpha peak. Since alpha oscillations have been linked to feedback communication, we suggest that the concurrent impairment in resting-state and stimulus-evoked alpha oscillations indicates an altered interaction of top-down and bottom-up processing in the visual hierarchy, which likely contributes to incomplete behavioral recovery in individuals who had experienced transient congenital blindness.

## Introduction

Dense bilateral congenital cataracts can cause complete patterned visual deprivation in humans (Birch et al., 2009). Delayed treatment results in increasingly severe visual impairment the later patients undergo surgery, which has been attributed to altered neural development instead of predominantly peripheral eye abnormalities (Maurer & Hensch, 2012; Röder & Kekunnaya, 2021). Based on invasive electrophysiological recordings in cochlear-implanted deaf cats (Yusuf et al., 2021, 2022), it was hypothesized that delayed sensory experience following birth particularly affects the elaboration of top-down processing (Röder & Kekunnaya, 2021), which is crucial for modulation of bottom-up signals. This hypothesis is consistent with feedback processing relying on corticocortical pathways, which showed a longer developmental trajectory than feedforward pathways in the mammalian visual cortex (Batardière et al., 2002; Andreas Burkhalter, 1993; Ibrahim et al., 2021).

Consistent with impaired feedback processing in the absence of visual experience, electroencephalography (EEG) studies in individuals treated for congenital cataracts reported reduced alpha (8-14 Hz) oscillations during visual motion tasks (Bottari et al., 2016, 2018). Multiple lines of evidence have linked alpha oscillations in the human EEG to feedback control of neural excitation in the visual cortex (Jensen & Mazaheri, 2010; Klimesch et al., 2007). Similar to findings from cataract-reversed individuals, permanently congenitally blind individuals also demonstrated lower alpha activity during non-visual tasks (Kriegseis et al., 2006) and at rest (Kriegseis et al., 2006; Novikova, 1974).

A special role of alpha oscillations in vision has been suggested by two widely-observed phenomena: first, the human EEG features prominent alpha oscillations at rest, which are enhanced with eye closure (Adrian & Matthews, 1934). Second, visual stimulation in the alpha range caused greater entrainment (Thut et al., 2011) of neural oscillations than at any other frequencies (Başar et al., 1997; Herrmann, 2001). Alpha oscillations were interpreted as the “natural frequency” at which an internal oscillator can be particularly entrained by stimulation, resulting in an enhanced response (Notbohm et al., 2016; VanRullen, 2016). Preferred neural responses in the alpha range were further observed in the impulse response function of the visual system to unbiased visual white-noise (Childers & Perry, 1971; Lalor et al., 2007). VanRullen and MacDonald (2012) employed white-noise stimulation, which changed in luminance with equal power between 0-80 Hz. Occipital EEG responses were cross-correlated with the presented luminance values, revealing a peak in the alpha range of the cross-correlation spectrum. This phenomena purportedly reflected an increased response of the visual system, typically oscillating at rest in the alpha range, to the alpha frequencies in the stimuli (Notbohm et al., 2016; Vanrullen & MacDonald, 2012). A correlation was observed between the amplitudes and frequencies of the resting-state and evoked alpha peaks, suggesting that both phenomena were linked. Based on such a relationship, and given the reduction of alpha oscillations in congenital blindness at rest (Novikova, 1974), we expected to find a reduction of both stimulus-evoked and resting-state alpha oscillations in cataract-reversed individuals.

Alternate accounts, however, interpreted evoked alpha oscillations as the linear addition of bottom-up activity from each stimulus, without reliance on resting-state alpha oscillations (Capilla et al., 2011; Keitel et al., 2019). Under this account, alpha reduction in cataract-reversed individuals might be specifically linked to visual processing, possibly as a consequence of impaired extrastriate processing (Sourav et al., 2018).

We adapted the EEG paradigm from VanRullen and MacDonald (2012) to elicit stimulus-evoked oscillations and unbiasedly determine the response properties of the visual cortex in sight-recovery individuals who underwent transient congenital blindness. Further, resting-state EEG was recorded with both eyes opened and eyes closed. Thirteen cataract-reversed individuals who had been born blind due to bilateral dense cataracts were compared to 13 age-matched, normally-sighted controls. We observed reduced alpha activity in the stimulus-evoked impulse response in cataract-reversed individuals, which was linked to lower visual acuity. The same individuals demonstrated reduced resting-state alpha oscillations. We interpreted these findings as evidence of compromised development of recurrent processing of the visual environment, resulting in poor vision.

## Methods

### Participants

#### We tested two groups of participants

The first group consisted of individuals who had been deprived of patterned vision at birth due to delayed treatment for dense bilateral congenital cataracts (CC individuals; n = 13, 3 female, Mean Age = 23.1 years, SD = 10.9 years, Range = 10.9 – 43.5 years) (Table 1). This group was recruited at the LV Prasad Eye Institute (LVPEI) by ophthalmologists and optometrists. Diagnosis of these patients was based on the following combination of criteria: the presence of dense bilateral congenital cataracts at birth, occlusion of the fundus, nystagmus, a diagnosis of dense bilateral congenital cataracts in immediate family members, and a visual acuity of counting fingers at 1 m or less prior to surgery, barring cases of absorbed lenses. Absorbed lenses occur specifically in individuals with dense congenital cataracts (Ehrlich, 1948). Participants with absorbed lenses were diagnosed based on the morphology of the lens, anterior capsule wrinkling and plaque or thickness of stroma. All 13 participants lacked stereovision, and 9 of 13 participants were implanted with intraocular lenses (Mean Visual Acuity = 0.85, SD = 0.40, Range = 0.17-1.78, all measured in logMAR units). Participants wore their optical corrections for the duration of the experiment. Maternal rubella was excluded as a cause of congenital cataracts in this group, explicitly excluded in the medical files of 11 participants, and not noted in the medical records of 2 participants (Supplementary Table S9).

**Table 1:** Demographic and clinical information of the participants with a history of dense bilateral congenital cataracts (CC). NA indicates patient’s data for the field were not available. FFL: Fixating and Following Light; CF: Counting Fingers; PL: Perceiving Light. Duration of visual deprivation was calculated by subtracting the date of birth from the date of surgery on the first eye. Time since surgery was calculated by subtracting the date of surgery in the first eye from the date of testing. Visual acuity reported was on the date of testing and measured using the Freiburg Vision Test (FrACT) (Bach, 2007).

|  | COMORBIDITIES |  |  | GENDER | VISUAL ACUITY PRE-SURGERY |  | DURATION OF VISUAL DEPRIVATION (YEARS) | TIME SINCE SURGERY (YEARS) | VISUAL ACUITY ON DATE TESTED (LOGMAR) | FAMILY HISTORY | INTRA-OCULAR LENS |
| --- | --- | --- | --- | --- | --- | --- | --- | --- | --- | --- | --- |
|  | ABSORBED LENSES | STRABISMUS | NYSTAGMUS |  | OD | OS |  |  |  |  |  |
| 1 | No | Yes | Yes | Male | FFL - | FFL + | 0.2 | 16.8 | 0.17 | No | Yes |
| 2 | Yes | Yes | Yes | Male | 1.18 | 1 | 20.8 | 22.7 | 0.9 | Yes | Yes |
| 3 | Yes | Yes | Yes | Male | 1.48 | 1.77 | 15.6 | 3.1 | 0.9 | No | No |
| 4 | Yes | NA | Yes | Male | CF at 1.5 m | CF at 3m | 7.0 | 8.2 | 0.62 | No | Yes |
| 5 | No | Yes | Yes | Male | NA | NA | 14.0 | 18.4 | 0.88 | Yes | Yes |
| 6 | No | Yes | Yes | Male | NA | NA | 6.0 | 17.9 | 0.78 | Yes | Yes |
| 7 | No | Yes | Yes | Male | PL+ | PL+ | 0.8 | 12.4 | 0.54 | No | No |
| 8 | Yes | Yes | Yes | Male | 1.2 | 1.3 | 16.4 | 1.9 | 0.66 | Yes | Yes |
| 9 | No | Yes | Yes | Male | NA | NA | 5.0 | 5.9 | 0.93 | Yes | No |
| 10 | No | Yes | Yes | Female | NA | NA | 5.0 | 11.0 | 0.5 | Yes | Yes |
| 11 | Yes | Yes | Yes | Female | 1.77 | 1.77 | 15.0 | 0.9 | 1.78 | Yes | Yes |
| 12 | No | NA | Yes | Male | NA | NA | 6.0 | 30.9 | 1.34 | Yes | No |
| 13 | Yes | Yes | Yes | Female | 1.48 | 1.48 | 31.4 | 7.4 | 1.04 | Yes | Yes |

All but two participants in this group were operated on after at least 1 year of patterned visual deprivation prior to surgery (Mean duration of blindness = 11 years, Range = 0.2 – 31.4 years). One participant received cataract removal surgery at the age of 3 months (Table 1). All participants were tested at least 11 months after receiving cataract removal surgery (Mean time since surgery = 12 years, Range = 0.9 – 30.9 years) to exclude acute effects and to allow for extensive visual experience.

The second group was recruited from the local area of the city of Hyderabad and consisted of normally sighted individuals (SC, n = 13, 2 female, Mean Age = 25.2 years, SD = 9.4 years, Range = 12.1 – 41.8 years). The two groups did not differ in mean age (t(24) = 0.52, p = 0.606).

All participants were tested at the LVPEI in English, Hindi or Telugu. None had a history of genetic, neurological or cognitive disorders. All participants (and their parents or legal guardians in case of minors) gave written informed consent. The study was approved by the Local Ethics Commission of the LVPEI, Hyderabad, India, as well as the ethics board of the Faculty of Psychology and Human Movement, University of Hamburg (UHH, Hamburg, Germany). Participants were compensated for costs associated with participation in this study including travel and accommodation, and children were additionally given a small gift.

### Procedure

Our experiment was adapted from VanRullen & MacDonald (2012) and modified for use in visually impaired individuals and children.

Stimuli were presented with a Dell laptop on a Dell 22-inch LCD monitor with a refresh rate of 60 Hz. Stimuli were created with MATLAB r2018b (The MathWorks, Inc., Natick, MA) and Psychtoolbox 3 (Brainard, 1997; Kleiner et al., 2007).

A session started with the recoding of resting-state EEG. Participants were asked to sit as still as possible during the recording, and either to fixate on a blank, dark screen (eyes open condition, EO) or to keep their eyes closed (eyes closed condition, EC). The order of these two conditions was randomized across participants, and each condition lasted for at least 3 minutes.

During the task, data were collected in a darkened room. In consecutive experimental trials, participants watched a circle at the center of a black screen. The visual angle subtended by the circle was 17 degrees. This circle changed in luminance between values ranging from 0 to 1. Luminance values changed with equal power at all temporal frequencies between 0-30 Hz, thus rendering each trial a white-noise luminance sequence (Figure 1, Supplementary Material S1), (Luo et al., 2021; Vanrullen & MacDonald, 2012). Unique, randomly generated white-noise sequences were presented for every trial and participant. Luminance values were gamma corrected to ensure a linear luminance output (monitor gamma value = 2.041). Participants were asked to perform 100 trials of a duration of 6.25 s each, divided into blocks of 10 trials.

**Figure 1:**
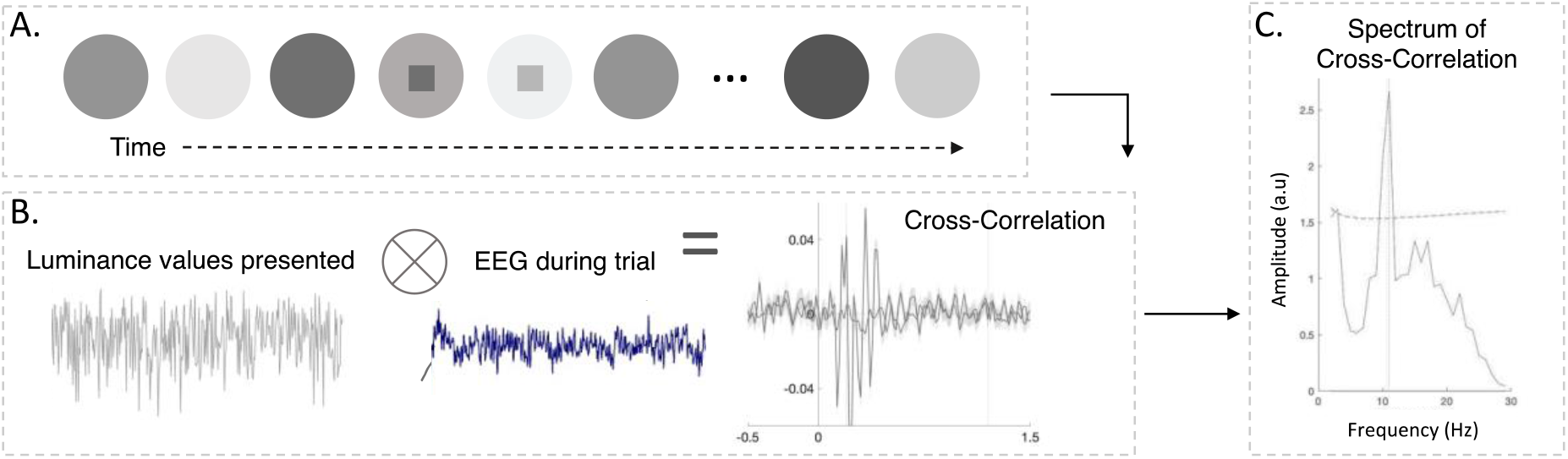
**A**. Schematic representation of the stimuli during a trial, and subsequent data analysis. Participants watched a circle that randomly changed in luminance. **B**. For each trial the EEG response was cross-correlated with the luminance value presented on that trial. **C**. The frequency spectrum of this cross-correlation analysis was calculated across all the trials. The peak frequency and amplitude of the cross-correlation spectrum was determined and used as a dependent variable.

In 10% of trials, a single square subtending a visual angle of 6 degrees appeared at the circle’s center. The luminance of this square was scaled to 0.9 times the luminance value of the circle. The target square would appear at the circle’s center for a randomly selected time, excluding the first 50 and last 50 frames. For the duration of its appearance, this square changed luminance at the same frequency as the circle. Participants were instructed to watch for the target square. At the end of every trial, participants verbally indicated whether or not they saw a square on that trial (Figure 1). The experimenter recorded the response (maximum response time = 10 s) (Supplementary Material S6). No feedback was provided on accuracy during the time of testing. At the end of every block, participants were asked whether they would like to take a break or continue with the task. Some participants terminated the experiment early; four participants (2 CC, 2 SC) performed 80 trials, and 1 CC participant performed 70 trials.

Prior to the beginning of the experiment, all participants performed 10 practice trials with a target appearing on 30% of them.

The EEG was recorded using Ag/AgCl electrodes attached according to the 10/20 system (Homan, Herman, & Purdy, 1987) to an elastic cap (EASYCAP GmbH, Herrsching, Germany). We recorded 32 channel EEG using a BrainAmp amplifier, with a bandwidth of 0.01–200 Hz and sampling rate of 5 kHz (http://www.brainproducts.com/). All scalp recordings were performed against a left ear lobe reference.

After the EEG recording, visual acuity was measured binocularly for every participant on the day of testing using the Freiburg Vision Test or FrACT (https://michaelbach.de/fract/, Bach, 2007). Visual acuity is reported as the logarithmic of the mean angle of resolution (logMAR), wherein higher values indicate worse vision (Elliott, 2016).

### Data analysis

#### Visual Stimulation EEG Analysis

The EEGLab toolbox in MATLAB 2018b was used for data analyses (Delorme & Makeig, 2004). First, datasets were filtered using a Hamming windowed sinc FIR filter, with a high-pass cutoff of 1 Hz and a low-pass cutoff of 45 Hz. The resulting data were down-sampled to 60 Hz (antialiasing filtering performed by EEGLab’s *pop_resample* function) to match the luminance stimulation rate. The data were divided into 6.25 s long epochs corresponding to each trial. Subsequently, baseline removal was conducted by subtracting the mean activity across the length of an epoch from every data point. After baseline removal, epochs with signals exceeding a threshold of ±120 μV were rejected in order to exclude artifacts.

Based on the existing literature (Lozano-Soldevilla & VanRullen, 2016, 2019; VanRullen, 2016; VanRullen & MacDonald, 2012) and pilot testing with 73 electrodes at University of Hamburg (see Supplementary Material S2), we analyzed data exclusively from the two occipital electrodes, O1 and O2. Recordings were referenced to the left earlobe. For each trial, the luminance values presented from 0.5 s to 5.75 s after trial onset were cross-correlated with the corresponding EEG time points during that trial. The initial and final portions were excluded from the analysis in order to eliminate the transients due to stimulus onset and offset (VanRullen & MacDonald, 2012). The average cross-correlation value across trials was computed across all visual stimulation epochs for every participant. As a control, we additionally calculated the cross-correlation of each EEG epoch with a luminance sequence that was presented on a different, randomly selected, trial. This control analysis was carried out to ensure that the evoked alpha response was specific to the stimuli presented and not an artifact of any kind of flickering stimulation (VanRullen & MacDonald, 2012).

Next, the amplitude spectrum of each participant’s cross-correlation function was calculated both for O1 and O2, for the delays between 0.2 s and 1.2 s. These delays were chosen as in VanRullen and MacDonald (2012). Using the *pwelch* function in MATLAB, we obtained the power spectral density of the cross-correlation (window length = 60 samples, overlap = 0, spectrum resolution = 1 Hz) (VanRullen & MacDonald, 2012).

Prior to identifying peaks in the spectrum of the cross-correlation function, we removed the aperiodic (1/f) component of this spectrum for each subject. Note that this analysis was not performed by VanRullen and MacDonald (2012). However, as has been suggested in several recent studies, we applied this correction to ensure that potential between-group differences in oscillations were not driven by differences in aperiodic activity (Donoghue, Haller, et al., 2020; Schaworonkow & Voytek, 2021). First, we fit the 1/f distribution function to the frequency spectrum of each participant. The 1/f distribution was fit to the normalized spectrum converted to logarithmic scale (range = 1-20 Hz) (Donoghue, Dominguez, et al., 2020; Gyurkovics et al., 2021; Schaworonkow & Voytek, 2021). We excluded the alpha range (8 – 14 Hz) for this fit, to avoid biasing the results (Donoghue et al., 2021; Schaworonkow & Voytek, 2021; Voytek et al., 2015; Waschke et al., 2017). This 1/f fit gave us a value of the slope, an overall intercept value corresponding to the broadband power of all frequencies, and a fit error for the cross-correlation spectrum of every participant. On subtracting the fitted 1/f spectrum from the original spectrum in logarithmic scale, we obtained the corrected cross-correlation spectrum for each subject between 1-30 Hz.

We used MATLAB’s *findpeaks* function to identify peaks between 7-14 Hz. Two criteria were used to define above-threshold peaks, in order to set quantitative criteria for whether an evoked alpha peak existed at all. First, the peak had to be higher than one standard deviation of the fit error, obtained from the 1/f fit of the cross-correlation spectrum. Second, peaks had to be at least 1 Hz (i.e. equal to the resolution of the spectrum) in width. Peak identification was performed individually for O1 and O2. Subsequently, for every subject, the peak frequency and amplitude were averaged across O1 and O2.

Finally, 1/f corrected spectra were averaged across O1 and O2 in order to obtain a mean cross-correlation spectrum for each subject.

#### Resting-State EEG Analysis

Resting-state data were preprocessed identically to the visual stimulation EEG data. We filtered the 3-minute-long resting-state recordings using a Hamming windowed sinc FIR filter (High and Low Cutoffs: 1-45 Hz). Next, we divided the recording into epochs of 1 s, and rejected epochs with signals exceeding ±120 μV. Finally, we calculated the power spectral density of the preprocessed EEG data using the *pwelch* function (window length = 1000 samples, overlap = 0).

We followed an identical procedure as the one described above for the cross-correlation spectrum to obtain peaks in the alpha range of the resting-state spectrum.

#### Experimental Design and Statistical Analysis

We hypothesized that stimulus-evoked occipital alpha activity in CC individuals was reduced compared to SC individuals. To test this hypothesis, first, the mean amplitude of the cross-correlation spectrum in the alpha range (8-14 Hz, averaged across O1 and O2) was compared between the two groups with a t-test. Second, a chi-square test was employed to test the likelihood that CC vs SC individuals presented an evoked alpha response. Third, for the subgroups of individuals who demonstrated above-threshold evoked alpha responses in their cross-correlation spectra, t-tests were conducted in order to compare the peak frequency and peak amplitude between CC and SC individuals.

Mean resting-state alpha activity was compared between groups. The average amplitude of the resting-state spectrum between 8-14 Hz for every subject (averaged across O1 and O2) was derived, and a group (2 levels: CC, SC) by condition (2 levels: EO, EC) ANOVA was performed on these average alpha amplitude values. Additionally, the likelihood of presenting an above-threshold resting-state alpha peak was compared between the two groups using Chi-squared tests. This was done separately for the EO and EC conditions. Finally, for the subgroup of individuals with above-threshold resting-state alpha peaks in the SC and CC groups, group-by-condition ANOVAs were performed in order to compare the peak frequency and peak amplitude values between these subgroups of CC and matched SC individuals.

As in VanRullen and MacDonald (2012), we tested for the presence of a correlation between the peak alpha frequency and amplitude values of the cross-correlation spectra, and the peak alpha frequency and amplitude in the resting-state (EC) spectra, respectively. These correlations were tested for the individuals who demonstrated an above-threshold alpha peak in the cross-correlation and resting-state (EC) spectra across both the CC and SC groups.

To determine whether vision loss history might predict the presence vs absence of alpha activity, CC participants were categorized based on whether or not they presented an above-threshold peak in the evoked alpha response, and compared on visual acuity, time since surgery, duration of blindness and age. Identical analyses were conducted comparing CC individuals categorized based on whether they presented above-threshold resting-state alpha activity, separately for the eyes closed and eyes opened conditions. Due to differing and small sizes of the subgroups of CC individuals, non-parametric testing was used to compare the subgroups via the Wilcoxon Rank Sum Test.

Anonymized data and materials will be made available upon reasonable request to the corresponding author, under stipulations of applicable law including, but not limited to the General Data Protection Regulation (EU 2016/679). This experiment was not pre-registered.

## Results

### 1. Reduced amplitude of stimulus-evoked alpha activity in congenital cataract-reversed individuals

First, we tested the cross-correlation of the EEG response with the corresponding white-noise luminance sequence (the average cross-correlation from each group is depicted in Figure 2A,B). As seen in Figure 2C, the average spectral density of this cross-correlation showed a peak in the alpha range for both the SC and the CC group. However, the mean power in the alpha range (8-14 Hz) was significantly reduced in the CC compared to the SC group (t(24) = −2.3, p = 0.028) (Figure 2C, 2D). While all SC participants displayed an above-threshold (see Methods) peak in the alpha range, only 8 of 13 CC participants did; the likelihood of an individual demonstrating an evoked alpha peak was significantly higher in the SC than in the CC group (Chi squared test: χ^2^ = 6.2, p = 0.013) (Figure 2C):

**Figure 2:**
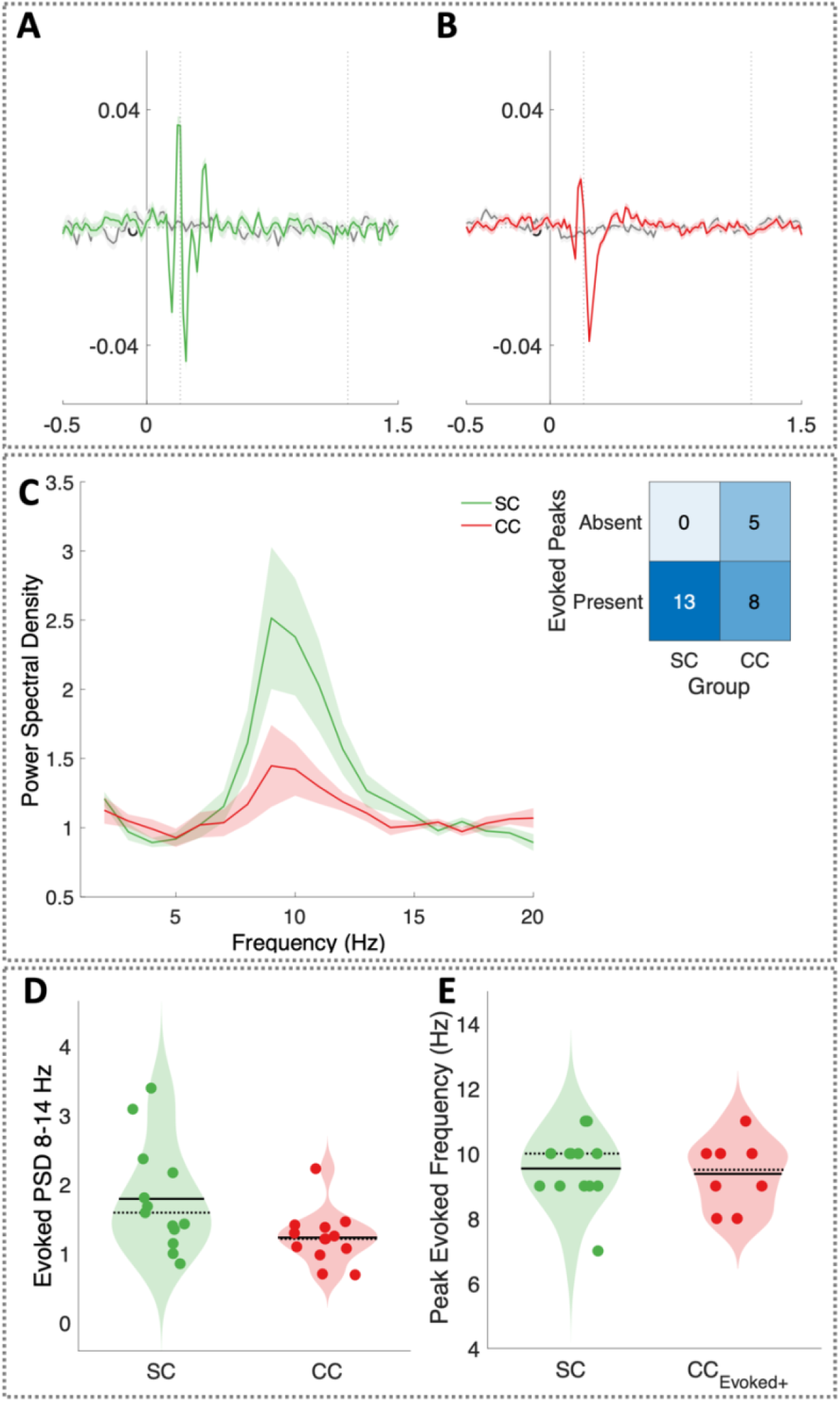
Stimulus-evoked alpha activity (in arbitrary units, a.u.) in congenital cataract-reversed (CC) and normally sighted (SC) individuals. **A,B**. The cross-correlation of the EEG response with the corresponding luminance values presented. Average cross-correlation response from participants of the A. SC (green) and the B. CC (red) group, respectively. The average cross-correlation across all trials is plotted in colour. The grey line represents the average cross-correlation of the EEG response on every trial with a randomly selected, mismatched luminance sequence. The shaded region represents the standard error of mean. The dotted lines mark the section of the cross-correlation used to calculate the spectrum. **C**. The average spectrum of cross-correlation functions across all CC (red) and SC (green) subjects. The shaded region represents standard error of the mean. Inset table listing the number of individuals in each group with and without an above-threshold evoked alpha peak. **D**. Violin plots displaying the average evoked alpha amplitude (averaged between 8-14 Hz) in SC (green) and CC (red) individuals. Solid lines indicate the mean values and dotted lines indicate median values of the average evoked alpha amplitude in the SC and CC group. Individual subjects have been horizontally jittered for a better view of overlapping data points. **E**. Violin plots displaying the peak frequency distributions of SC and CC_Evoked_+ individuals, the subgroup of CC individuals who presented an above-threshold evoked alpha peak.

For the subgroup of 8 CC individuals with an above-threshold evoked alpha peak (henceforth referred to as CC_Evoked+_), peak frequency values did not differ from those observed for the SC group (t(19) = 0.3, p = 0.73) (Figure 2E). Moreover, the peak evoked alpha amplitude did not significantly differ between the CC_Evoked+_ subgroup and the SC group (t(19) = 1.35, p = 0.19) (Supplementary Material S3).

### 2. Reduced amplitude of resting-state alpha activity in congenital cataract-reversed individuals

To test the effects of transient early visual deprivation on resting-state alpha oscillations, we compared the alpha range of the resting-state spectra between CC and SC individuals. CC individuals showed an overall reduction of the average alpha power (8-14 Hz) compared to SC individuals, in both the eyes open (EO) and the eyes closed (EC) conditions (main effect of group: F(1,48) = 9.6, p = 0.003) (Figure 3A,C). Across groups, alpha amplitude was significantly lower in the eyes open (EO) than in the eyes closed (EC) condition (main effect of condition: F(1,48) = 11.5, p = 0.001). The reduction of alpha power from the EC to the EO condition was indistinguishable between the CC and SC group (group-by-condition interaction F(1,48) = 0.96, p = 0.333) (Figure 3A,C).

**Figure 3:**
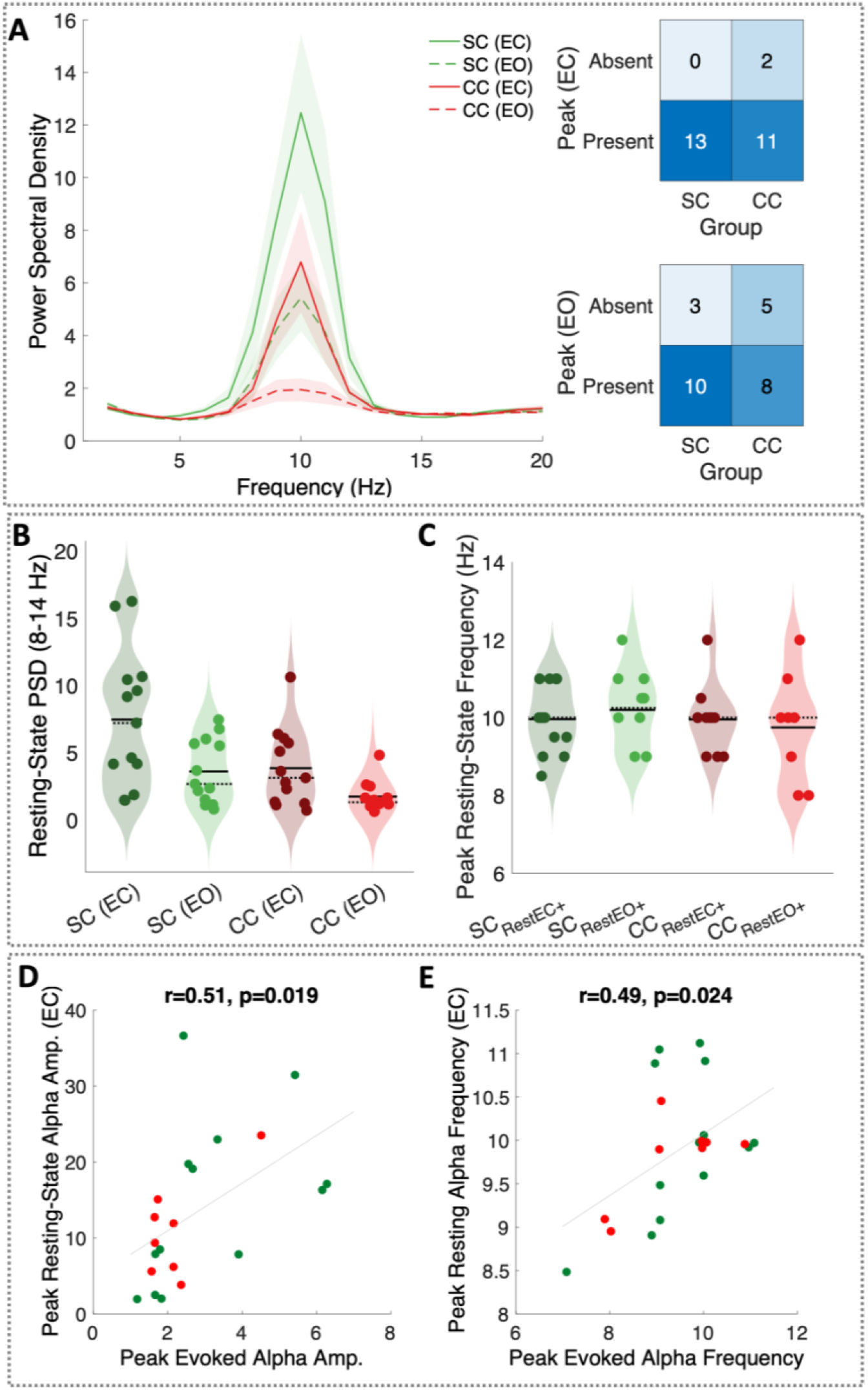
Resting-state alpha oscillations of congenital cataract-reversed (CC) and normally sighted control (SC) individuals. **A**. Mean resting-state spectra with eyes opened and eyes closed, averaged across all SC (green) and all CC (red) individuals. Shaded regions represent the standard error of the mean. Inset tables listing the number of individuals in each group with and without an above threshold alpha peak, for the eyes open and the eyes closed condition. **B**. Violin plots illustrate the average (8-14 Hz) resting-state alpha amplitude distributions for SC and CC individuals. Solid lines indicate the mean values and dotted line indicate median values of the average evoked alpha amplitude in the SC and CC group. **C**. Violin plots illustrate the peak alpha frequency distributions for the subgroup of SC and CC individuals with an above-threshold resting-state alpha peak in either the eyes open (SC_RestEO+_, CC_RestEO+_) or eyes closed (SC_RestEC+_, CC_RestEC+_) condition. **D,E**. Pearson correlation between stimulus-evoked and resting-state alpha oscillations. Correlations between values of D. amplitude and E. frequency of the resting-state and evoked alpha peaks across CC_Evoked+_ (red) and SC (green) individuals.

The likelihood of demonstrating an above-threshold (see Methods) peak in the alpha range of the resting-state spectrum did not differ between the SC and CC groups, neither in the EO (Chi squared test: χ^2^ = 0.72, p = 0.395) nor in the EC condition (Chi squared test: χ^2^ = 2.17, p = 0.141) (Figure 3B).

Within the subgroups of CC and SC individuals with above-threshold resting-state alpha peaks (SC group: n = 13 in the EC condition, 10 in the EO condition, henceforth referred to as SC_RestEC+_ and SC_RestEO+_ respectively; CC group: n = 11 in the EC condition, 8 in the EO condition, henceforth referred to as CC_RestEC+_ and CC_RestEO+_ respectively), the peak frequency value did not differ between the EO and EC conditions (main effect of condition: F(1,34) = 0.06, p = 0.81). Moreover, the subgroups of SC_RestEO+_, SC_RestEC+_, CC_RestEC+_ and CC_RestEO+_ individuals did not differ in peak frequency (main effect of group: F(1,34) = 2.3, p = 0.13, group-by-condition interaction: F(1,34) = 1.08, p = 0.31) (Figure 3D).

Finally, there was a marginal reduction of the peak resting-state alpha amplitude in the CC_RestEC+_ and CC_RestEO+_ subgroups compared to the SC_RestEO+_ and SC_RestEC+_ subgroups (main effect of group: F(1,34) = 3.07, p = 0.089), and the expected increase of peak alpha amplitude with eye closure across subgroups (main effect of condition: F(1,34) = 8.17, p = 0.007, group-by-condition interaction: F(1,34) = 0.05, p = 0.821) (Supplementary Material S4).

Across the SC_Evoked+_ and CC_Evoked+_ subgroups (n = 21), there was a significant positive correlation between the peak resting-state alpha frequency and the peak evoked alpha frequency (r = 0.49, p = 0.024) and between the peak resting-state alpha amplitude and the peak evoked alpha amplitude (r = 0.51, p = 0.019) (Figure 3E,F).

### 3. Concurrent reduction of stimulus-evoked and resting-state alpha power in congenital cataract-reversed individuals

As expected from the mechanistic account described in VanRullen and MacDonald (2012), the subgroup of CC_Evoked-_ individuals, with no stimulus-evoked alpha peak (N=5), had a significantly lower resting-state alpha amplitude than CC_Evoked+_ individuals, who demonstrated such a peak (N=8) (main effect of subgroup: F(1,22) = 14.3, p = 0.001) (Figure 4A). Resting-state alpha power increased with eye closure in both subgroups (main effect of condition: F(1,22) = 7.3, p = 0.013). However, the magnitude of this effect was larger in CC_Evoked+_ than in CC_Evoked-_ individuals (subgroup-by-condition interaction F(1,22) = 4.61, p = 0.043; Figure 4B).

**Figure 4:**
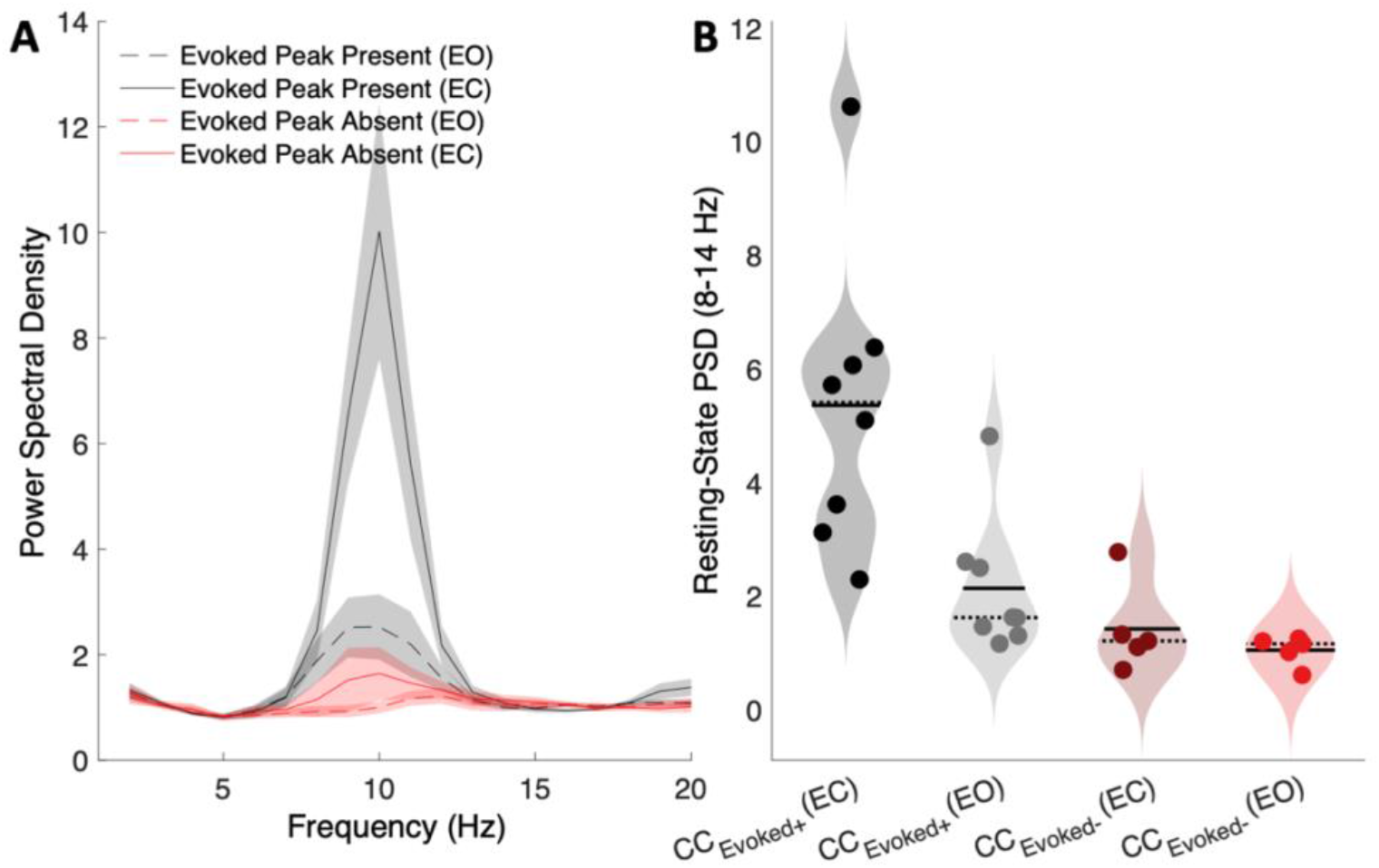
**A**. Mean resting-state spectra plotted for the eyes opened and eyes closed conditions in the subgroup of cataract-reversed individuals with (CC_Evoked+_, n = 8, black) and without (CC_Evoked-_, n = 5, red) an above-threshold evoked alpha peak. Shaded region represents the standard error of mean. **B**. Violin plots depict the mean resting-state alpha amplitude of CC_Evoked+_ and CC_Evoked-_ individuals.

Post-hoc testing revealed lower alpha power in the EC condition in CC_Evoked-_ than CC_Evoked+_ individuals (t(11) = 3.3, p = 0.007), while this difference was marginally significant for the EO condition (t(11) = 1.96, p = 0.076).

### 4. Lower visual acuity in cataract-reversed individuals without an above-threshold evoked alpha response

Visual acuity was significantly better, that is, logMAR values were lower, in CC_Evoked+_ (n =8) than CC_Evoked-_ (n = 5) individuals (Wilcoxon rank-sum test: z = 2.2, p = 0.0286) (Figure 5). When we compared visual acuity in CC individuals with vs without an above-threshold resting state peak, there was no difference in visual acuity between CC_RestEC+_ (n = 11) vs CC_RestEC_-individuals (n = 2) in the eyes closed condition (Wilcoxon rank-sum test: z = 1.09, p = 0.277), or between CC_RestEO+_ (n = 7) vs CC_RestEO_-individuals (n = 6) in the eyes open condition (Wilcoxon rank-sum test: z = 0.14, p = 0.886) (Figure 5).There was no subgroup difference in chronological age, time since surgery or duration of blindness between CC_Evoked+_ and CC_Evoked-_ individuals (Supplementary Material Figure S7).

**Figure 5:**
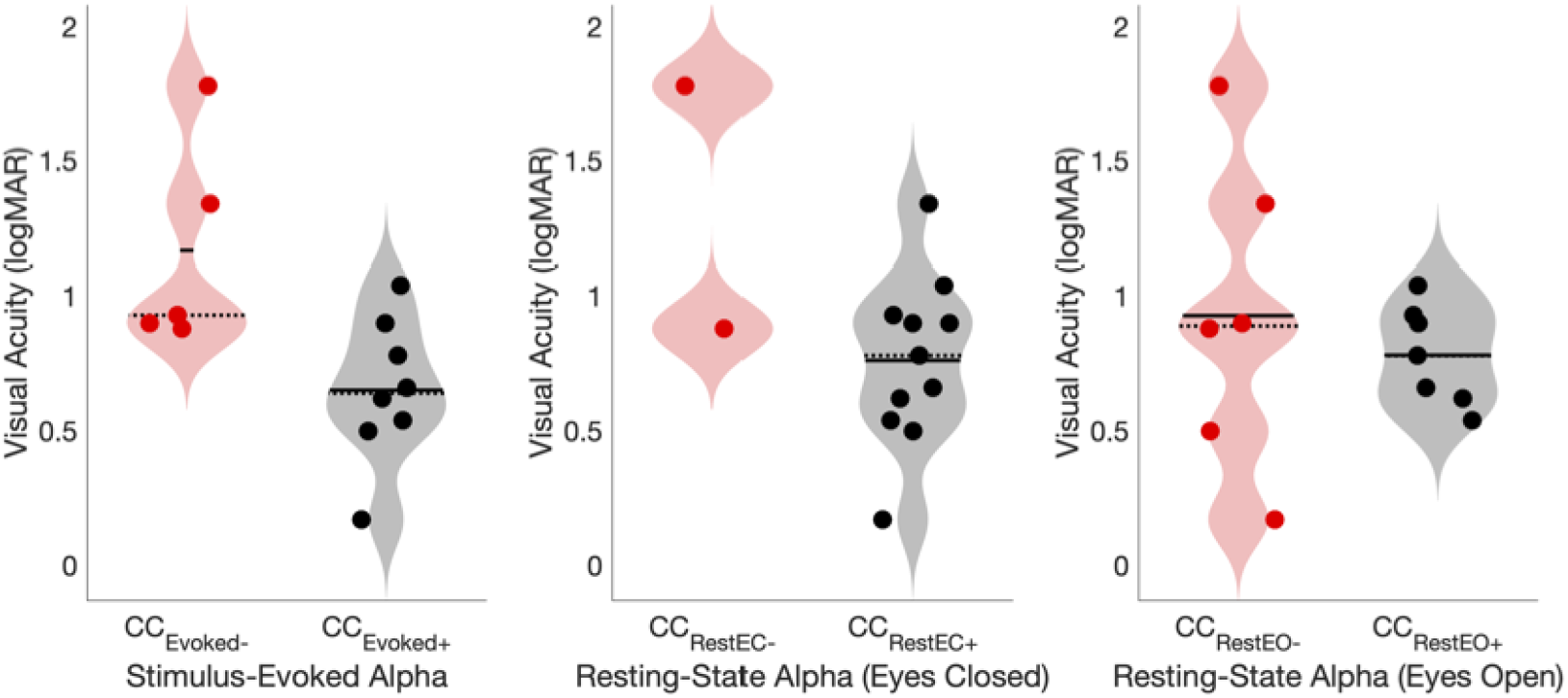
Violin plots depict the visual acuity in logMAR units in the CC group divided into subgroups with (black) and without (red) an above-threshold peak in the stimulus-evoked (CC_Evoked+_, CC_Evoked-_), resting-state (CC_RestEC+_, CC_RestEC-_, eyes closed) and resting-state (CC_RestEO+_, CC_RestEO-_, eyes open) conditions.

## Discussion

The present study investigated stimulus-evoked and resting-state alpha (8-14 Hz) oscillations in individuals who had experienced congenital blindness for an average of 11 years, due to delayed surgery for dense bilateral congenital cataracts. We used the association between stimulus-evoked and resting-state alpha as a proxy to assess bottom-up and top-down processing in the visual system and how it depends on early visual experience in humans (see Yusuf et al., 2022, for a non-human model of auditory deprivation). Stimulus-evoked oscillations were assessed with the visual impulse response to white noise. In normally sighted individuals (SC) we replicated the main results from VanRullen and MacDonald (2012): we observed the characteristic prominent stimulus-evoked alpha response (Başar et al., 1997; Childers & Perry, 1971; Lalor et al., 2007; Vanrullen & MacDonald, 2012). Individuals treated for congenital cataracts (CC) demonstrated a lower amplitude of both stimulus-evoked and resting-state alpha oscillations, as compared to the SC group. Stimulus-evoked and resting-state alpha oscillations were correlated in amplitude and peak frequency across CC and SC individuals. Finally, visual acuity was worse in CC individuals who did not demonstrate an above-threshold evoked alpha peak. The present study provides electrophysiological evidence for the crucial role of early visual experience in the joint emergence of stimulus-evoked and resting-state alpha oscillations. These findings are compatible with the idea that the elaboration of top-down pathways, which functionally shape upstream sensory areas to process the visual environment, is experience-dependent; and likely linked to a sensitive period in human brain development.

Multiple lines of evidence have linked alpha oscillations to feedback communication in the visual system, including results from non-human animal studies (Maier et al., 2010; Van Kerkoerle et al., 2014) invasive recordings from humans at rest (Halgren et al., 2019), non-invasive EEG recordings (Händel et al., 2011; Jensen & Mazaheri, 2010; Klimesch et al., 2007; Worden et al., 2000) and transcranial magnetic stimulation studies in humans (Riddle et al., 2020; Sauseng et al., 2011; Zanto et al., 2011). Stimulus-evoked alpha activity has been proposed to result from entrainment of the cortical generators of resting-state oscillations (Başar et al., 1997; Herrmann, 2001; Herrmann et al., 2016; Notbohm et al., 2016; Zoefel et al., 2018) and thus, likely reflects an interaction of top-down and bottom-up signals. Evidence for an enhanced propensity of the visual cortex to generate alpha oscillations was demonstrated by assessing the impulse response to visual white noise (Vanrullen & MacDonald, 2012). Visual white noise comprises equal stimulation at all frequencies; nevertheless, visual circuits entrain predominantly in the alpha range. The correlation between the peak frequency and amplitude of stimulus-evoked and resting-state alpha oscillations, also observed in the present study, is compatible with a link between stimulus-evoked and resting state alpha oscillations (Vanrullen & MacDonald, 2012; Zoefel et al., 2018).

Within the entrainment account of stimulus-evoked alpha oscillations, we speculate that jointly reduced stimulus-evoked and resting-state alpha oscillations might reflect an altered interaction of top-down and bottom-up processing streams, or recurrent processing (Yusuf et al., 2022), in the visual cortex. Non-human animal models have provided three lines of evidence for this idea. First, the lack of visual experience caused a flattening of the cortical hierarchy in macaque visual cortex (Magrou et al., 2018). In sighted macaques, the laminar ratio of supragranular to the sum of supragranular and infragranular neurons decreases from lower to higher visual regions, and is considered a quantitative measure of interareal distance in the cortical hierarchy (Markov et al., 2014). In enucleated macaques, this laminar ratio was altered, which might imply impaired interareal communication in the visual cortex (Magrou et al., 2018).

Second, feedback interareal projections in the visual cortex have been shown to be the subject of experience-dependent shaping to a greater extent than feedforward projections in macaques (Batardière et al., 2002), humans (Burkhalter et al., 1993) and rodents (Ibrahim et al., 2021). Thus, the connectivity necessary to iteratively adjust resting-state activity to optimize visual processing might be particularly vulnerable to the effects of early visual deprivation (Pezzulo et al., 2021). Third, analogous observations of altered feedback processing after sensory restitution were made in electrophysiological recordings from the auditory cortex of congenitally deaf cats, stimulated via cochlear implants. Feedback coupling from secondary to primary auditory cortex, measured by phase consistency, was particularly lowered by congenital auditory deprivation (Yusuf et al., 2021). Functional coupling between infragranular and supragranular layers in A1 was reduced in cochlear-implanted compared to normally hearing cats (Yusuf et al., 2022) suggesting an impairment of the interaction of feedback and feedforward processing streams. Similar to the present work, the authors suggested that the lack of sensory experience at birth interrupts the sculpturing of feedback connectivity in sensory cortex, resulting in impaired orchestration of bottom-up and top-down processing pathways (Kral et al., 2017; Yusuf et al., 2021, 2022).

Reduced alpha entrainment would be consistent with lower visual capabilities in CC individuals. Previous studies have found that an alpha entrainment improves sub-threshold detection (Spaak et al., 2014) and distractor suppression (Wiesman & Wilson, 2019). Accordingly, in our study the individuals with reversed congenital cataract who featured a significant stimulus-evoked alpha peak were those with better visual acuity outcomes. Interestingly, a study which optogenetically inactivated top-down signals from marmoset V2 to V1 found increased receptive field sizes in V1 (Nurminen et al., 2018), which implies lower visual acuity. Assuming that lower alpha activity is associated with impaired feedback tuning of early visual cortex, it would be justified to conclude that a deficit of this mechanism contributes to the lower visual acuity as found in the present study and repeatedly documented in CC individuals. Further, our interpretation of impaired recurrent processing would be consistent with other visual deficits reported in CC individuals, including visual feature binding (McKyton et al., 2015; Putzar et al., 2007), coherent motion processing (Hadad et al., 2012; Rajendran et al., 2020) and face identity processing (Le Grand et al., 2001; Putzar et al., 2010). Higher visual functions require the integration of outputs of multiple neural circuits within and across visual areas, and thus, functional interareal connectivity. For example, deactivating top-down connectivity from the mouse homologous area V2 to V1 resulted in a reduced firing to illusory contours (Pak et al., 2020), reminiscent of a similar deficit in CC individuals (McKyton et al., 2015; Putzar et al., 2007). In addition to the present results, another electrophysiological study in CC individuals found evidence for impaired interareal communication. Pitchaimuthu et al. (2021) recorded steady-state visual evoked potentials to visual stimuli simultaneously changing two features - flickering and simultaneously moving horizontally. Such combined stimulation typically results in intermodulation frequency responses in the EEG indicate the integration of input across multiple visual areas (Kim et al., 2011). Intermodulation frequency responses were absent in CC individuals indication independent evidence for an impaired integration across visual regions (Pitchaimuthu et al., 2021).

The present findings are compatible with prospective developmental results from ferrets and humans. Invasive recordings in the ferret visual cortex demonstrated that the similarity between stimulus-driven and resting-state activity increased during early development (Berkes et al., 2011). Moreover, studies in humans have documented a protracted developmental trajectory for resting-state alpha oscillations into adolescence (Cellier et al., 2021; Marshall et al., 2002). Rare results on stimulus-evoked alpha responses in children suggested maturation beyond early childhood (Kolev et al., 1994). Therefore, we hypothesize that the congenital blindness prevented the development of characteristic resting-state activity, which might prepare visual circuits to efficiently response to visual input. As a consequence, stimulus-evoked processing might be less efficient, resulting in worse visual behavior.

Testing the integrity of visual circuits in humans who were treated for bilateral dense cataracts could be considered analogous to the prevalent approach in non-human animals, wherein experimentally manipulated visual deprivation is used to study the effects of experience on brain development. Limitations of the human model have been discussed (Röder & Kekunnaya, 2021), and here we acknowledge some challenges for the present study. Humans who had experienced a period of visual deprivation longer than about 8 weeks following birth typically suffer from nystagmus (Supplementary Table S9), and therefore differences in involuntary eye movements between the CC and SC groups are confounded with our measurements. However, nystagmus might not explain the present results, as all CC participants suffered from nystagmus, but eight out of thirteen showed a stimulus-evoked alpha peak. Moreover, differences in resting-state EEG are unlikely to be linked to fixation abilities.

Further, non-invasively recorded alpha oscillations cannot unambiguously being equated with top-down activity. Testing for a modulation of alpha power with top-down cues in CC individuals would be necessary to further investigate whether top-down modulation of restored bottom-up input is affected by congenital visual deprivation in humans. Initial evidence suggested that despite an overall reduction of activity in CC compared to SC individuals in visual motion areas (hMT), CC individuals displayed increased hMT activity if the motion was task relevant (Guerreiro et al., 2022). In agreement with the present findings, these results suggest that top-down control of upstream visual areas is less stimulus-specific and precise, but not absent in individuals who recovered vision after congenital blindness

## Acknowledgments

We are grateful to D. Balasubramanian and the LV Prasad Eye Institute for supporting the study. We thank Kabilan Pitchaimuthu, Suddha Sourav and Prativa Regmi for helping with the data acquisition. The study was funded by the German Research Foundation (DFG Ro 2625/10-1 and SFB 936-178316478-B11) to Brigitte Röder. Rashi Pant was supported by a PhD student fellowship from the Hector Fellow Academy gGmbH.

