## Supplementary Material for "Stimulus-evoked and resting-state alpha oscillations show a linked dependence on patterned visual experience for development"

### S1. Schematic of white-noise luminance spectrum generation

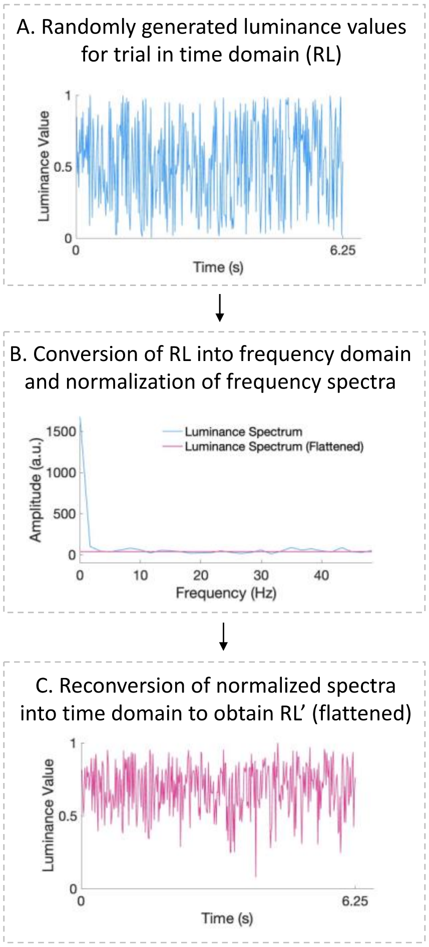


*Figure S1. Schematic of generating white-noise sequences of luminance values. A. An example of the random luminance sequence initially generated (blue). B. Original frequency decomposition of randomly generated sequence (blue) and the normalized flattened spectrum of the same sequence (pink). C. Reconversion of flattened spectrum from the previous step into luminance values corresponding to it. These luminance values are presented to the participant.*

### S2. Pilot testing

Pilot testing of the visual task described in Methods was conducted with two normally sighted participants (Ages: 26 years, 21 years) in a darkened, electrically and acoustically shielded room at the University of Hamburg, (Germany). We used a 73 electrode EEG setup and a monitor with a refresh rate of 120 Hz. Using the same data analysis procedure as described in Methods, we replicated results from VanRullen and MacDonald (2012). Both pilot participants demonstrated a peak in the alpha range of the cross-correlation response spectra. Like in VanRullen and MacDonald (2012), this response was most expressed at the occipital electrodes (O1 and O2).

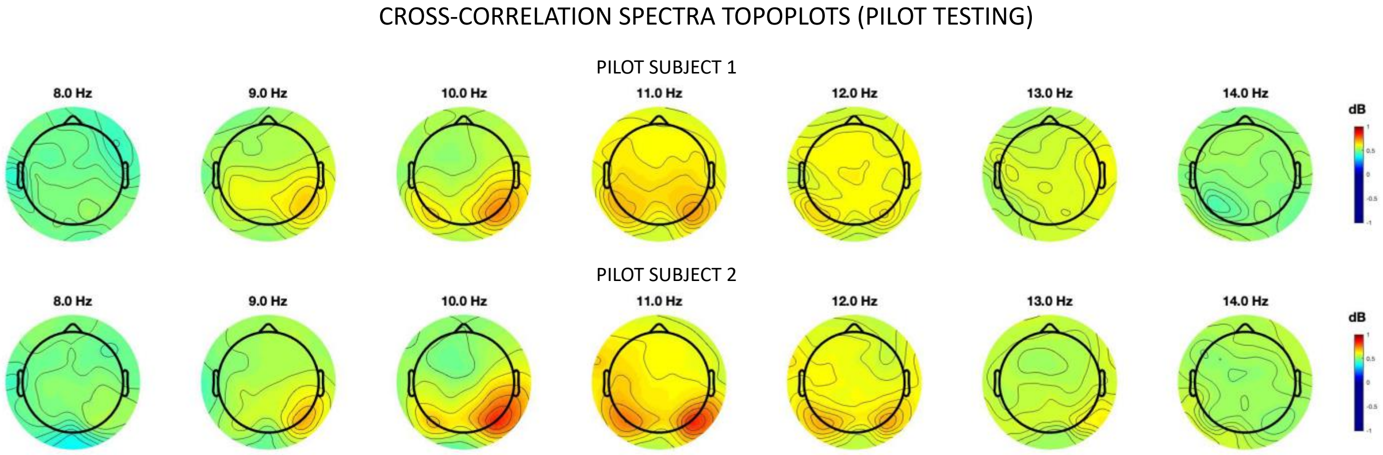


*Figure S2: Topographic representation displaying the power (dB) of the cross-correlation spectra of the EEG response and the luminance changes. Cross-correlation spectra from two normally sighted pilot participants are represented as a heatmap on the scalp, with higher values (red) corresponding to higher power at frequencies between 8-14 Hz.*

**S3. Peak amplitude of the stimulus-evoked spectrum**

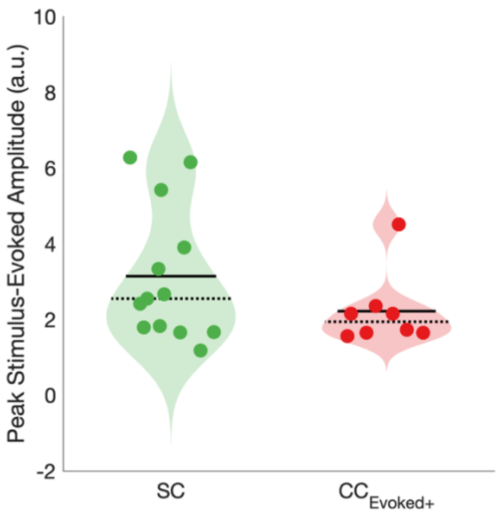


*Figure S3: Violin plot showing the distribution of the stimulus-evoked alpha amplitude at the peak frequency of normally sighted (SC) individuals and cataract-reversed individuals who presented above-threshold evoked alpha peaks ($CC_{Evoked+}$).*

### S4. Peak alpha amplitude of the resting-state spectrum

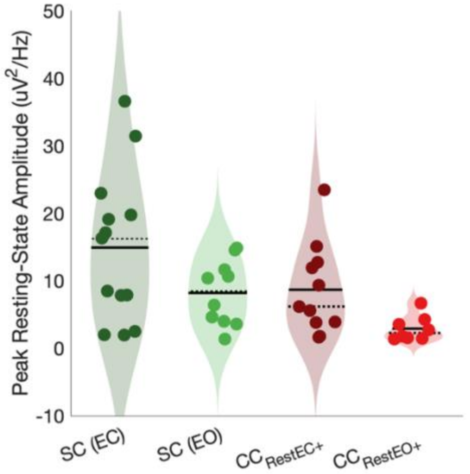


*Figure S4: Violin plot showing the distribution of the peak resting-state alpha amplitude in normally sighted (SC) and cataract-reversed (CC) individuals who presented above-threshold resting-state alpha peaks in the eyes open (RestEO+) and eyes closed (RestEC+) conditions.*

### S5. Correlation analysis of demographic data with peak stimulus-evoked alpha frequency and amplitude

The effect of chronological age on peak evoked alpha frequency and average evoked alpha amplitude across both the CC and SC groups was tested. Chronological age predicted neither frequency nor amplitude of evoked alpha activity across groups (Figure S5). Next, within the CC group, we tested whether duration of visual deprivation, time since surgery and visual acuity affected the peak evoked alpha frequency and average evoked alpha amplitude. None of these demographic and clinical factors were correlated with average amplitude or peak frequency of the evoked alpha oscillations (Figure S5).

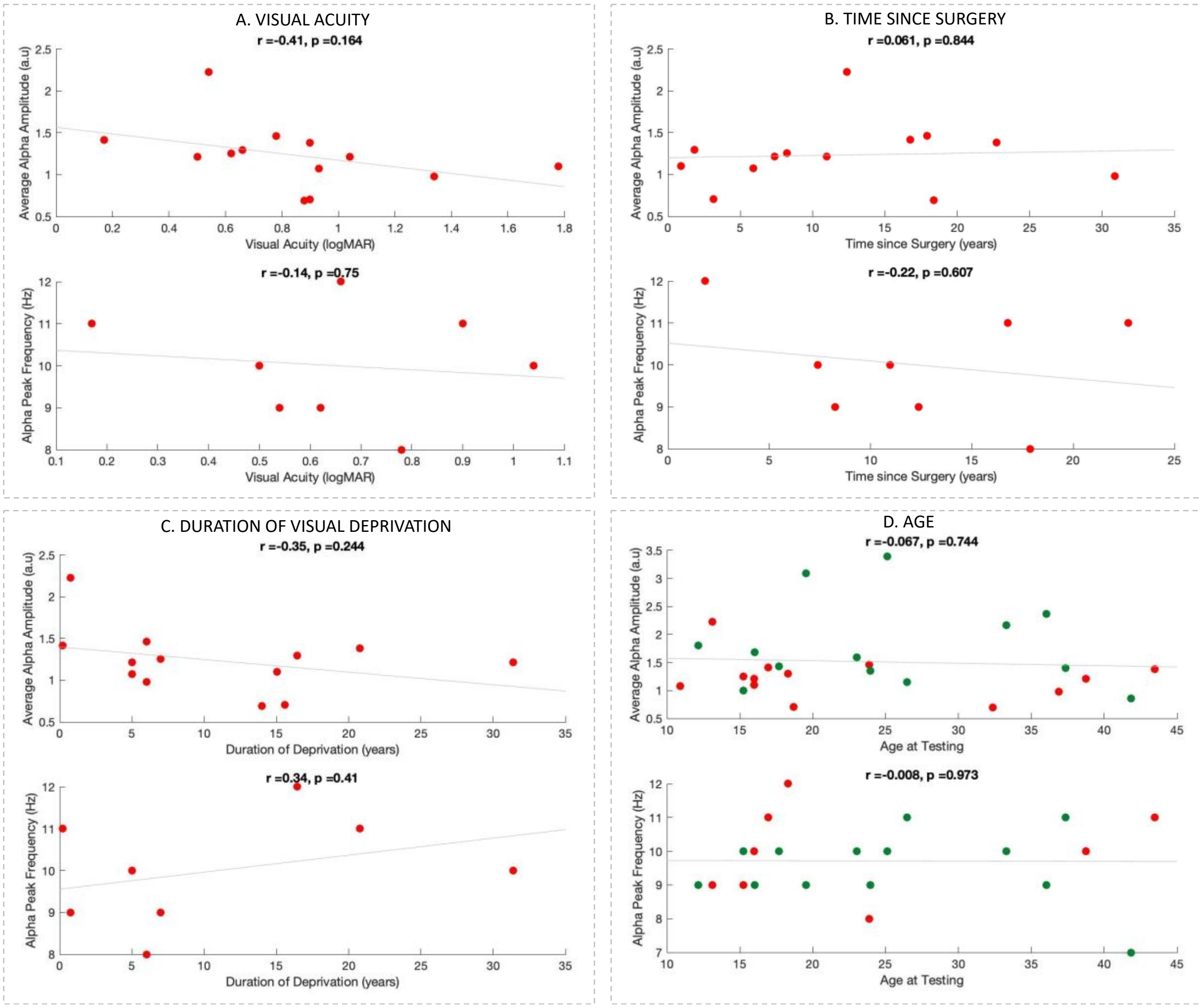


*Figure S5: Correlations between the average stimulus-evoked alpha amplitude and peak evoked alpha frequency and demographic factors within the CC and SC groups. Correlations of the average alpha amplitude (top panel) and the alpha peak frequency (lower panel) with A. visual acuity in the CC group (measured using FrACT on the date of testing), B. time since surgery in the CC group (calculated by subtracting the date of surgery in the first eye from the date of testing), C. duration of visual deprivation (calculated by subtracting date of birth from date of surgery in the first eye) in the CC group and D. chronological age at testing in the CC (red) and SC (green) group*

### S6. Behavioral data

We tested behavioral performance on the target detection task across all trials. Participants were asked to indicate at the end of trial whether or not they saw a target on that trial. Accuracy was defined as the ratio of correctly indicated rejections and hits to the total number of trials performed by the participants. Accuracy did not differ between groups ($t(12) = -1.012$, $p = 0.331$).

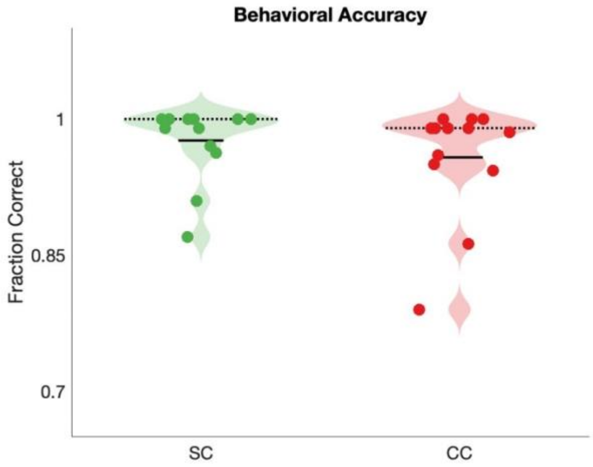


*Figure S6: Violin plot showing the distribution of the accuracy on target detection in cataract-reversed (CC) and age-matched sighted control (SC) individuals.*

**S7. Demographic data of cataract-reversed individuals with an above-threshold evoked alpha response present vs absent**

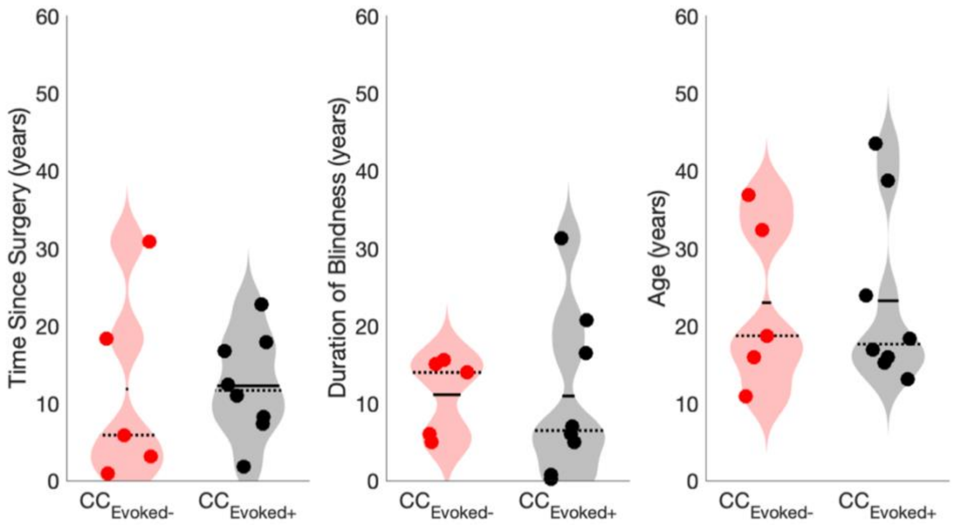


*Figure S7 Violin plots showing time since surgery, duration of blindness and chronological age of cataract-reversed individuals with ($CC_{Evoked+}$ black, present) and without ($CC_{Evoked-}$, red, absent) an above-threshold stimulus-evoked alpha peak.*

### S8. Topoplots of stimulus-evoked alpha activity in SC and CC individuals

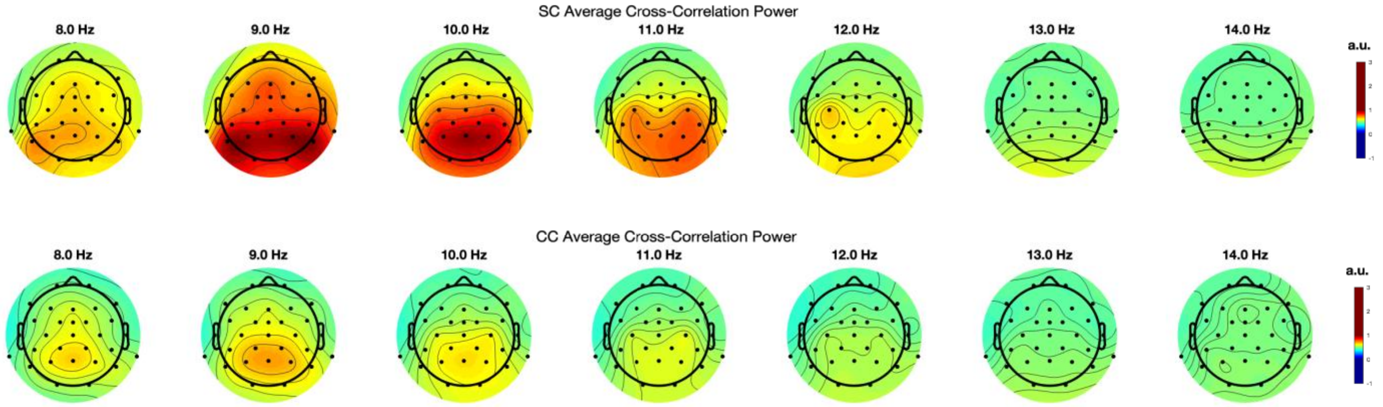


*Figure S8. Topographic representation displaying the average power (a.u.) of the cross-correlation spectra of the EEG response and the luminance changes normally sighted (SC) participants (top) and cataract-reversed individuals (CC). Power is represented as a heatmap on the scalp, with higher values (red) corresponding to higher power of the cross-correlation at frequencies between 8-14 Hz.*

### S9. Extended participant aetiology

**(Excel Spreadsheet)**

*Table S9***:** *Extended clinical information of the participants with a history of dense bilateral congenital cataracts (CC). NA indicates patient's data for the field were not available. FFL: Fixating and Following Light; CF: Counting Fingers; PL: Perceiving Light. Duration of visual deprivation was calculated by subtracting the date of birth from the date of surgery on the first eye. Time since surgery was calculated by subtracting the date of surgery in the first eye from the date of testing. Visual acuity reported was on the date of testing and measured using the Freiburg Vision Test (FrACT)* (Bach, 2007). *Fundus photography was conducted for participant numbers 8,9 and 13.*

### S10. Relationship between visual acuity improvement and alpha oscillations

In order to assess whether CC patients' stimulus-evoked alpha power showed a clear benefit of surgery with significantly improved visual acuity values, we tested the correlation between the difference in logMAR values Post-Pre surgery and stimulus-evoked alpha power in the 8-14 Hz range. Note that negative Post-Pre surgery logMAR values indicate greater improvement in vision. Further, for participants with Pre-Sx visual acuities in the CF range, we used plausible extensions to the FrACT scale in logMAR units as described by the authors (Lange et al., 2009; Schulze-Bonsel et al., 2006). Improvement in visual acuity did not significantly predict the amplitude of stimulus-evoked alpha activity (r = -0.62, p = 0.103).

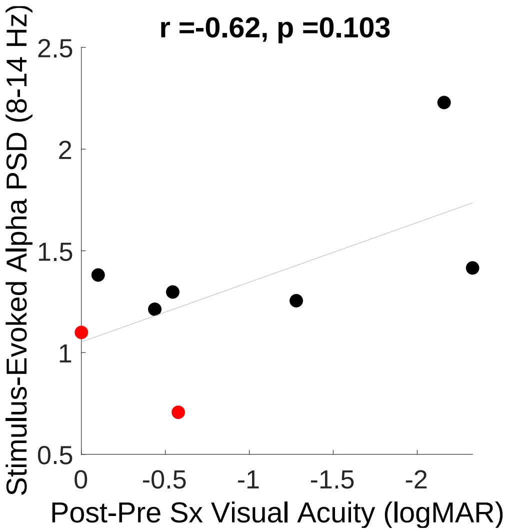


*Figure S10. Correlation between stimulus-evoked alpha power and change/improvement in visual acuity after cataract removal. Note that the x-axis is inverted in direction to indicate greater improvement in vision.*
